# The striatal compartments, striosome and matrix, are embedded in largely distinct resting state functional networks

**DOI:** 10.1101/2024.12.13.628392

**Authors:** Alishba Sadiq, Adrian T. Funk, Jeff L. Waugh

## Abstract

The striatum is divided into two interdigitated tissue compartments, the striosome and matrix. These compartments exhibit distinct anatomical, neurochemical, and pharmacological characteristics and have separable roles in motor and mood functions. Little is known about the functions of these compartments in humans. While compartment-specific roles in neuropsychiatric diseases have been hypothesized, they have yet to be directly tested. Investigating compartment-specific functions is crucial for understanding the symptoms produced by striatal injury, and to elucidating the roles of each compartment in healthy human skills and behaviors. We mapped the functional networks of striosome and matrix in humans *in vivo*. We utilized a diverse cohort of 674 healthy adults, derived from the Human Connectome Project, including all subjects with complete diffusion and functional MRI data and excluding subjects with substance use disorders. We identified striatal voxels with striosome-like and matrix-like structural connectivity using probabilistic diffusion tractography. We then investigated resting state functional connectivity (rsFC) using these compartment-like voxels as seeds. We found widespread differences in rsFC between striosome-like and matrix-like seeds (*p* < 0.05, FWE corrected for multiple comparisons), suggesting that striosome and matrix occupy distinct functional networks. Slightly shifting seed voxel locations (<4 mm) eliminated these rsFC differences, underscoring the anatomic precision of these networks. Striosome-seeded networks exhibited ipsilateral dominance; matrix-seeded networks had contralateral dominance. Next, we assessed compartment-specific engagement with the triple-network model (default mode, salience, and frontoparietal networks). Striosome-like voxels dominated rsFC with the default mode network bilaterally. The anterior insula (a primary node in the salience network) had higher rsFC with striosome-like voxels. The inferior and middle frontal cortices (primary nodes, frontoparietal network) had stronger rsFC with matrix-like voxels on the left, and striosome-like voxels on the right. Since striosome-like and matrix-like voxels occupy highly segregated rsFC networks, striosome-selective injury may produce different motor, cognitive, and behavioral symptoms than matrix-selective injury. Moreover, compartment-specific rsFC abnormalities may be identifiable before disease-related structural injuries are evident. Localizing rsFC differences provides an anatomic substrate for understanding how the tissue-level organization of the striatum underpins complex brain networks, and how compartment-specific injury may contribute to the symptoms of specific neuropsychiatric disorders.

## 1 Introduction

The striatum serves as the primary subcortical connection hub, playing an essential role in a diverse array of motor and cognitive functions (Haber, Kim et al. 2006; Jung, Jang et al. 2014; Graybiel and Grafton 2015). Abnormalities in both the structure and function of the striatum have been associated with a range of neurological conditions, including Parkinson disease (Albin, Young and Penney 1989), Huntington disease (Rosenblatt and Leroi 2000), autism spectrum disorder (ASD) (Schuetze, Park et al. 2016), and schizophrenia (Chakravarty, Rapoport et al. 2015). The human striatum is composed primarily of medium spiny neurons (MSNs) that develop in two spatially segregated tissue compartments, the striosome and matrix. Compartment-specific functions demonstrated in animals suggest that compartment-selective injury could lead to distinct symptoms in human diseases. Over a dozen human diseases exhibit compartment-selective injury or a pattern of symptoms that suggests a compartment-selective injury (Crittenden and Graybiel 2011; Marecek, Krajca et al. 2024). Recent studies extended the understanding of compartment-specific disease mechanisms beyond the Crittenden and Graybiel review (2011), suggesting that additional disorders, including ASD (Kuo and Liu 2020), substance use disorders (Salinas, Davis et al. 2016), and certain neuropsychiatric conditions such as mood disorders in early Huntington disease (Waldvogel, Thu et al. 2012; Hedreen, Berretta and White III 2024) or anxiety disorder (Karunakaran, Amemori et al. 2021), also exhibit differential effects on the striosome and matrix compartments, indicating that the number of diseases with compartment-selective pathologies is likely higher than previously proposed. To the best of our knowledge, these hypothesized compartment-symptom associations have not been tested in living humans: tissue must be *ex vivo* (for fixation and immunohistochemical staining) to distinguish the striatal compartments, limiting the exploration of human diseases with striatal pathologies. Although the striosome and matrix were recognized in brain tissue four decades ago, the lack of tools to identify the compartments in living organisms has made it difficult to study striosome- and matrix-specific functions.

The striosome and matrix compartments exhibit distinct pharmacologic characteristics that suggest a basis for separate functional roles. The striosome is enriched with mu-opioid receptors (MORs) and has lower levels of calbindin, a neurochemical profile that aligns them with the modulation of dopamine-related signaling, crucial for emotional regulation and reward processing (McGregor, McKinsey et al. 2019). Limbic-related regions, such as the prelimbic cortex (PL), basolateral amygdala (Ragsdale Jr and Graybiel 1988; McGregor, McKinsey et al. 2019), and anterior cingulate (Eblen and Graybiel 1995) selectively project to the striosome. The striosome, but not the matrix, selectively inhibits the dopaminergic projection neurons of the substantia nigra pars compacta (SNc) (Crittenden, Tillberg et al. 2016; McGregor, McKinsey et al. 2019), emphasizing their involvement in reward-based learning and behavioral reinforcement. Eblen and Graybiel demonstrated in macaque that limbic cortices selectively project to the striosome, and that these projections are organized somatotopically within the striatum (Eblen and Graybiel 1995). Likewise, the striosome plays a crucial role in integrating inputs from these limbic regions, which are essential for regulating behavior and emotional responses. Plotkin and Prager identified a differential response to dopamine between striosome and matrix compartments, noting that dopamine release is modulated differently in each compartment, potentially influencing distinct behavioral outcomes, and contributing to the pathophysiology of neuropsychiatric disorders (Prager and Plotkin 2019). These findings suggest that the striosome may also be involved in the pathophysiology of neuropsychiatric disorders. These findings underscore that striosome function is closely linked to regulation of nigral dopamine signaling and coordination of activity among limbic regions, emphasizing the role of the striosome in emotional and motivational processing (Crittenden and Graybiel 2011).

In contrast, the matrix is characterized by higher calbindin and lower MOR expression, is selectively targeted by projections from somatomotor cortices (primary sensory cortex, primary motor cortex, supplementary motor area; Gerfen 1984; Donoghue and Herkenham 1986) and projects to the primary output nuclei of the striatum, the globus pallidus interna (Gimenez-Amaya and Graybiel 1990) and substantia nigra pars reticulata (Desban, Gauchy et al. 1989). This structural organization highlights the role of the matrix in motor control and sensorimotor integration, distinct from the striosome. Considering the compartments’ distinct inputs and projections, relatively segregated distributions within the striatum, opposing responses to dopaminergic regulation (Prager and Plotkin 2019), and selective susceptibility to metabolic injury (Burke and Baimbridge 1993), there are multiple anatomic bases to propose different functions for striosome and matrix. The differential functions of the striatal compartments are further evidenced by the striosome’s role in stereotypic behaviors (Lewis and Kim 2009), task engagement (Friedman, Homma et al. 2015), and reinforcement learning, particularly in encoding reward prediction errors (Barto 1995; Houk, Adams and Barto 1995) and valence discrimination (Friedman, Hueske et al. 2020). In contrast, the matrix compartment does not significantly influence these functions, suggesting a more specialized role for striosome in these aspects of behavior and learning (Graybiel and Matsushima 2023).

We previously demonstrated (Waugh, Hassan et al. 2022) that differential structural connectivity (probabilistic diffusion tractography) distinguishes striatal voxels with striosome-like and matrix-like patterns of structural connectivity. This method replicates the compartment-specific structural connectivity demonstrated through decades of injected tract tracer studies in animals (Funk, Hassan et al. 2023; Funk, Hassan and Waugh 2024), and is highly reliable in living humans, with a 0.14% test-retest error rate (Waugh, Hassan et al. 2022). These findings support the notion that the human striatum is anatomically organized into distinct compartments, each characterized by unique patterns of structural connectivity. Striatal compartmental organization is not merely structural, however, but also has functional implications (Donoghue and Herkenham 1986; Flaherty and Graybiel 1993; Eblen and Graybiel 1995). Specifically, these studies highlight how corticostriatal connectivity is directed primarily through one striatal compartment, suggesting a potential basis for distinct functional processing of biased corticostriate projections. Although compartment-specific functions are only partially explored in humans, research in animals provides evidence that striosome and matrix have distinct functional roles. For example, the striosome is specifically involved in decision-making under threat and in the formation of negative-valence memories (Saka and Graybiel 2003; Xiao, Deng et al. 2020; Nadel, Pawelko et al. 2021). These findings suggest that the striosome may play a primary role in processing emotionally salient information and guiding behavior in response to negative stimuli. However, despite these insights, a significant gap remains in our understanding of the compartment-specific functions of the striatum, particularly in humans. Moreover, it is unclear how each compartment contributes to more complex or nuanced behaviors, and whether functional distinctions observed in animal models are directly applicable to human behaviors. Further research is needed to delineate the specific roles of striosome and matrix in human cognitive and emotional processing, such as how the compartments interact within larger neural networks.

The human brain achieves complex and contingent regulation of functions in part by modulating interacting and competing networks of connected regions (Menon 2011). Understanding cognitive and behavioral functions depends on the structure of large-scale brain networks (Bressler and Menon 2010). Among the many stable intrinsic brain networks, Menon (2011) proposed a ‘triple-network model’ that highlighted the interplay of activation and regulation among three fundamental neurocognitive networks: the default mode network (DMN), salience network (SN), and frontoparietal network (FPN). Further, selective functional abnormalities in the triple-network are characteristic of specific neuropsychiatric disorders (Menon 2011). Resting-state fMRI studies have consistently identified reduced DMN activity in Alzheimer disease (AD; Binnewijzend, Schoonheim et al. 2012; Zhong, Huang et al. 2014; Li, Yao et al. 2017), major depressive disorder (MDD; Wei, Qin et al. 2015), and autism spectrum disorder (ASD; Wang, Li and Niu 2021). Specific types of disruptions in triple-network connectivity are associated with particular neuropsychiatric disorders.

Gaining insight into how the striosome and matrix compartments regulate human brain networks can improve our understanding of the triple-network model and its effects on cognition and behavior. The present study highlights a novel assessment of resting state functional connectivity (rsFC) in the striosome-like and matrix-like compartments of the striatum in living humans and explores the influence of each compartment on both whole-brain networks and the triple-network. Our examination included a relatively large adult cohort, comprising 674 healthy individuals of both sexes and diverse racial backgrounds. To the best of our knowledge, this is the first study investigating the functional connectivity differences between the striatal compartments in human subjects. Our findings suggest that striosome-like and matrix-like voxels are embedded in largely distinct functional networks. Therefore, compartment-selective injury or maldevelopment may underlie the network derangements and specific symptoms of neurodevelopmental disorders that involve the striatum.

## 2 Materials and Methods

In Figure 1, we summarize the methodology for investigating compartment-specific rsFC in living humans. Briefly, we utilized differential structural connectivity to parcellate the striatum into voxels with striosome-like and matrix-like structural connectivity profiles, generating individualized striatal masks for each subject and hemisphere. These masks follow the spatial distribution, relative abundance, and extra-striate structural connectivity patterns identified through animal and human histology (Waugh, Hassan et al. 2022; Funk, Hassan et al. 2023; Funk, Hassan and Waugh 2024). We then used these compartment-specific masks as the seeds for rsFC analyses.

**Figure 1:**
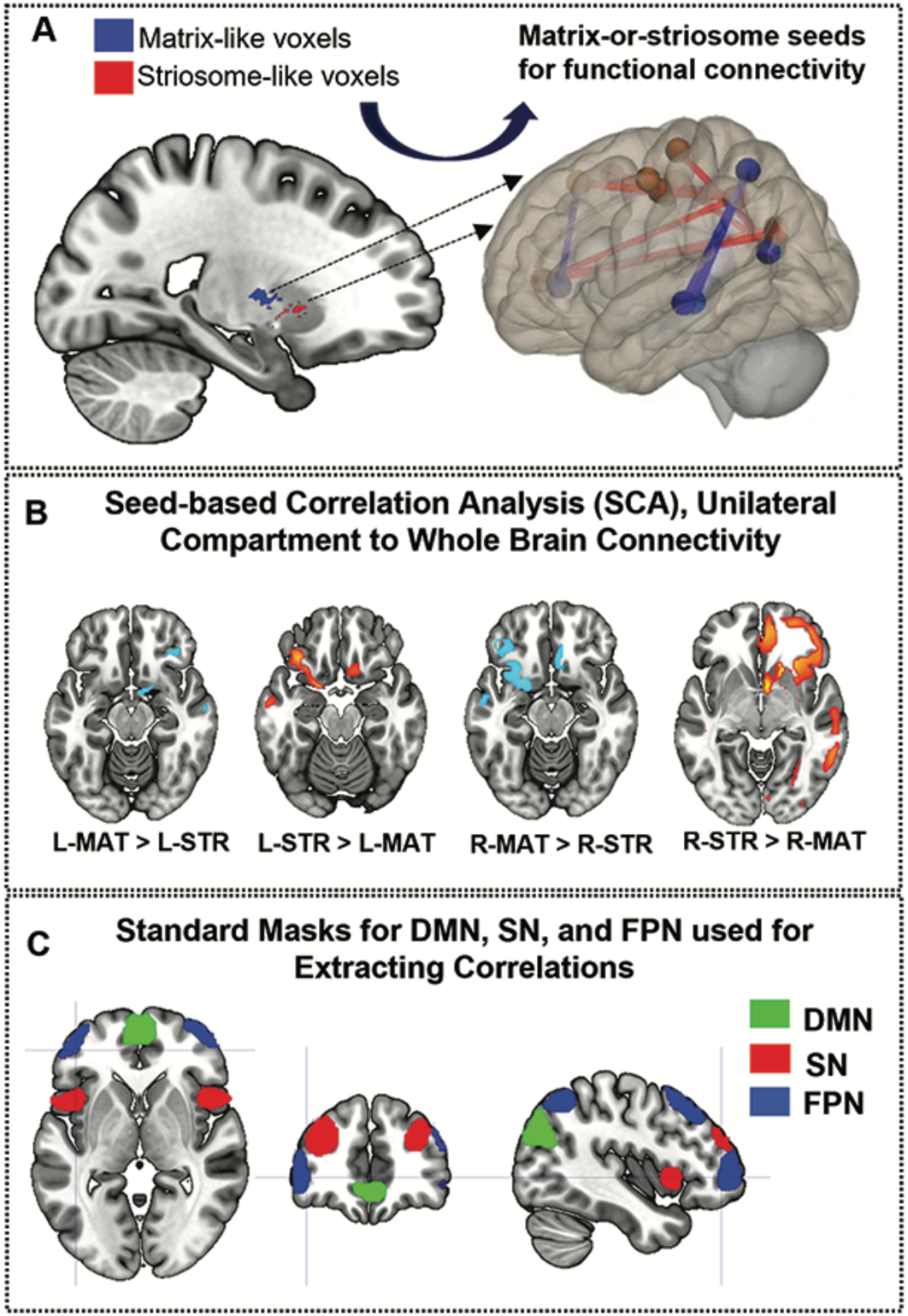
Framework for the investigation of compartment-specific differences in resting-state functional connectivity. (A) This sagittal section of the brain illustrates striosome-like (red) and matrix-like (blue) voxels, defined by differential structural connectivity. These voxels were used as seeds for functional connectivity analysis, identifying regions where matrix-like voxels show greater connectivity than striosome-like voxels or vice versa (B), with lighter-hued colors indicating stronger differences. (C) illustrates the nodes utilized to define the triple network: default mode network (DMN - green), salience network (SN - red), and frontoparietal network (FPN - blue). L-MAT: left matrix. L- STR: left striosome. R-MAT: right matrix. R-STR: right striosome.

### 2.1 Study Population

The data utilized in this study were sourced from the Human Connectome Project S1200 Release (Van Essen, Smith et al. 2013), which contained behavioral and MRI data from 1206 healthy young adults. We selected subjects that had full complements of diffusion MRI and resting state fMRI data, then eliminated subjects with a history of substance or alcohol use disorders (defined by DSM-5 criteria). We excluded subjects if they had any history of cocaine, hallucinogen, cannabis, nicotine, opiate, sedative, or stimulant drug use in their lifetime. We excluded subjects if they met the DSM criteria for Alcohol Abuse or Alcohol Dependence, or if they averaged ≥4 drinks per week over the year preceding their scan. This exclusion criteria produced a cohort of 674 healthy adults (mean age: 28.8 years, SD = 3.7 years). Three hundred and ninety-one participants were female; two hundred and eighty-three were male. All participants gave written informed consent (Van Essen, Ugurbil et al. 2012).

### 2.2 Acquisition of MRI data

All individuals were scanned at 3T using MRI sequences that were harmonized across multiple sites. The resting state fMRI data were collected across four runs, each lasting approximately 15 minutes. Two runs occurred within one session, and the other two occurred within a separate session. Within each session, oblique axial acquisitions alternated between phase encoding in a right-to-left (RL) direction for one run and in a left-to-right (LR) direction for the other run. Participants’ eyes were open and fixated on a crosshair. In this study, we employed the resting-state session 1 for both LR and RL acquisitions, concatenating 1200 timepoints for each session. Consequently, the resulting concatenated image comprises 2400 timepoints. Other acquisition parameters included: echo time (TE) = 33 ms, repetition time (TR) = 720 ms, flip angle = 52 deg; 72 slices; 2.0 mm isotropic voxels; acquisition time = 14 min and 33s. Diffusion tensor imaging (DTI) data for S1200 subjects was acquired at 1.25 mm isotropic resolution using 200 directions (14 B_0_ volumes, 186 volumes at non-colinear directions) with the following parameters: repetition time = 3.23 s; echo time = 0.0892.

### 2.3 Striatal Parcellation

We previously described a technique for parcellating the human striatum into compartments with striosome-like and matrix-like patterns of connectivity *in vivo* (Waugh,Hassan et al. 2022). We refer to parcellated striatal voxels as “striosome-like” or “matrix-like” due to the inferential nature of this method, and to remind readers that these voxels are not the equivalent of striosome and matrix tissue identified through immunohistochemistry. Striatal parcellation uses structural connectivity findings from decades of injected tract tracer investigations in mice, rats, cats, dogs, and non-human primates to identify regions with compartment-selective projections. We utilized five striosome-favoring regions, and five matrix-favoring regions, as competing targets (“bait”) for quantitative probabilistic tractography (connectivity-based parcellation), carried out in native diffusion space. The relative connectivity to these two groups of bait regions defined the compartment-like bias for each striatal voxel, mapping striosome-like and matrix-like probability distributions for each individual and hemisphere. We chose the most-biased voxels from each distribution to represent striosome and matrix, setting equal volume masks for each compartment. The summed volume of these highly biased masks was set to equal the uppermost 1.5 standard deviations above the volume of the striatal seed mask. We have demonstrated that this threshold is sufficient to recapitulate the relative abundance, spatial distribution, and selective connectivity of striosome and matrix that were demonstrated in histology (Waugh, Hassan et al. 2022; Funk, Hassan et al. 2023). In the current study we used these parcellated striatal voxels as seeds for rsFC with either whole-brain networks (excluding the striatum), or with nodes in the triple-network.

Striosome- and matrix-like voxels are uniquely located in each individual; while the striosome is enriched in rostral, medial, and ventral parts of the striatum (Graybiel and Ragsdale Jr 1978; Goldman-Rakic 1982; Donoghue and Herkenham 1986; Ragsdale Jr and Graybiel 1990; Desban, Kemel et al. 1993; Eblen and Graybiel 1995; Waugh, Hassan et al. 2022), at any particular location one may find striosome-like or matrix-like voxels. We measured the location of each striosome-like and matrix-like voxel to assess whether our parcellated voxels matched the compartment-specific location biases established in histology. For each subject and hemisphere, we identified the cartesian position (x, y, and z planes) of each compartment-like voxel. We then measured the within-plane and root-mean-square distances from that voxel to the centroid of the nucleus (caudate or putamen) in which the voxel resided.

We previously demonstrated that slight shifts in voxel location were sufficient to eliminate any compartment-like biases in structural connectivity, indicating that compartment-specific connectivity biases were specific for those precise locations, not for the “neighborhood” in which they reside (Funk, Hassan et al. 2023). We hypothesized that compartment-specific biases in functional connectivity would also be dependent on precise voxel location. We jittered the location of our compartment-like voxels by ±0-3 voxels, at random and independently in each plane. For the first 100 subjects in this cohort, we tested the cartesian position of each striosome- and matrix-like voxel before and after jitter to assure that while individual voxels shifted position, the mean location of these randomized masks had the same topographic organization as our compartment-specific masks. No voxels in the randomly shifted masks were reselected from either original compartment-like mask.

### 2.4 Analysis of Resting State Functional Connectivity (rsFC)

Functional connectivity measures were calculated between striatal seed masks (striosome-like, matrix-like, or randomized location) and all other non-striatal voxels in the brain. First, we computed the Pearson’s correlation coefficients between the time course of the residual BOLD and the time courses of all other voxels in the brain to estimate the first-level correlation maps of that time course. Correlation values were transformed to fisher’s z-score to normalize the distribution of data. The rsFC between each compartment-like seed and each component of the triple-network was evaluated using a series of network-specific masks from the connectivity (CONN) toolbox (Whitfield-Gabrieli and Nieto-Castanon 2012). We evaluated the three limbs of the triple-network: the default mode network (DMN), comprising the medial prefrontal cortex (mPFC), bilateral lateral parietal cortex, and posterior cingulate cortex (PCC, which also included parts of the precuneus); the salience network (SN), consisting of the dorsal anterior cingulate cortex (dACC), bilateral anterior insula (AI), bilateral rostral prefrontal cortex (PFC), and bilateral supramarginal gyrus (SMG); and the frontoparietal network (FPN), including the bilateral dorsolateral prefrontal cortex (dlPFC) and bilateral posterior parietal cortex (PPC). All 15 regions of interest (ROIs) were sourced from the CONN toolbox, which generated these target masks by independent component analysis (ICA) studies using the 497-subject Human Connectome Project (HCP) dataset (Van Essen, Ugurbil et al. 2012). By using publicly available ROIs, we minimized the possibility of bias that could result from manually selecting ICAs derived from our own data set, strengthening our external validation. Each ROI’s connectivity with striosome-like and matrix-like voxels was assessed independently, rather than performing a full-factorial analysis.

### 2.5 Compartment-Specific Bias in Functional Connectivity: N-1 Analysis

We previously demonstrated that this set of bait regions, which included the anterior insula, produced robust striatal parcellations (Waugh, Hassan et al. 2022). However, it is not possible to use a region to parcellate the striatal compartments and then accurately measure connectivity between the compartments and that same region. To avoid this issue when assessing connectivity with the SN (which included the anterior insula as a node), we performed a post-hoc validation in which we excluded the anterior insula from the set of bait regions and parcellated the striatum again using the remaining four striosome-favoring regions and original five matrix-favoring regions. We used this “leave one out” parcellation to seed rsFC only for measures that included the anterior insula, referred to here as an “N-1 analysis” (Funk, Hassan et al. 2023).

### 2.6 Statistical Analysis

First, we carried out whole-brain rsFC mapping. Statistical analyses were conducted on the correlation maps generated from striosome- and matrix-like seeds. Each subject contributed four maps, corresponding to connectivity dominated by four distinct seed regions: 1) left-matrix (L-MAT), 2) left-striosome (L-STR), 3) right-matrix (R-MAT), and 4) right-striosome (R-STR). Differences in correlation maps with the rest of the brain (excluding the striatum) were observed.

Next, we carried out rsFC mapping for the nodes of the DMN, SN, and FPN to assess compartment-specific alterations within the triple-network. These analyses were executed using SPM12 software (https://www.fil.ion.ucl.ac.uk/spm/software/spm12/). Paired t-tests were employed to compare the compartment-specific correlation maps (connectivity with striosome-like seeds vs. connectivity with matrix-like seeds) for each network-specific ROI. We modeled age and gender as covariates. We set a cluster-level threshold of p ≤ 0.05, corrected for the family-wise error (FWE) rate, indicating a significant difference in rsFC between connectivity seeded by striosome-like and matrix-like voxels.

## 3 Results

### 3.1 Relative Abundance and Location of Striosome-like and Matrix-like Voxels

Prior studies determined that the striosome and matrix comprise approximately 15% and 85% of striatal volume, respectively, in both animal and human histology (Johnston, Gerfen et al. 1990; Desban, Kemel et al. 1993; Holt, Graybiel and Saper 1997; Mikula, Parrish et al. 2009). We assessed the total number of highly biased (P>0.87) striosome-like and matrix-like voxels for each subject. Striosome-like voxels made up 5.9% of highly biased voxels, while matrix-like voxels made up 94.1%. Note that most striatal voxels have middling-bias (a blend of striosome- and matrix-like connectivity) and thus are not counted in this metric. These results align with histology, showing far more matrix-like than striosome-like voxels.

The relative intra-striate location of striosome and matrix is highly consistent across species (mouse, rat, cat, macaque, human), with striosome enriched in the rostral, ventral, and medial striatum and the matrix enriched in the caudal, dorsal, and lateral striatum (Graybiel and Ragsdale Jr 1978; Goldman-Rakic 1982; Donoghue and Herkenham 1986; Ragsdale Jr and Graybiel 1990; Desban, Kemel et al. 1993; Eblen and Graybiel 1995; Waugh, Hassan et al. 2022). Our voxel-by-voxel location analysis recapitulated this compartment-specific location bias in both hemispheres. In the caudate we found that matrix-like voxels were more lateral (0.88 mm, p=1.3×10^-78^), caudal (7.9 mm, p<1×10^-260^), and dorsal (8.2 mm, p<1×10^-260^) than striosome-like voxels. In the putamen we found that matrix-like voxels were shifted more lateral (2.4 mm, p<1×10^-260^), caudal (5.6 mm, p<1×10^−260^), and dorsal (6.3 mm, p<1×10^-260^) than striosome-like voxels. Striosome-like voxels in the caudate were 10.8 mm medio-rostro-ventral to the centroid (RMS distance), while matrix-like voxels were 11.5 mm latero-caudo-dorsal to the centroid. Striosome-like voxels in the putamen were 9.8 mm medio-rostro-ventral to the centroid, while matrix-like voxels were 7.5 mm latero-caudo-dorsal to the centroid.

### 3.2 Hemispheric Differences in Striosome-like and Matrix-like Functional Connectivity

We observed significant differences in whole-brain rsFC between striosome-like and matrix-like seeds, including notable compartment-specific differences in ipsilateral vs. contralateral connectivity (Figures 2, 3). The R-STR exhibited the highest hemispheric bias, with 92.8% of its significant voxels connected within the ipsilateral hemisphere. Conversely, connectivity with the R-MAT was dominated by contralateral connectivity, with 90.9% of its significant voxels in the left hemisphere.

**Figure 2:**
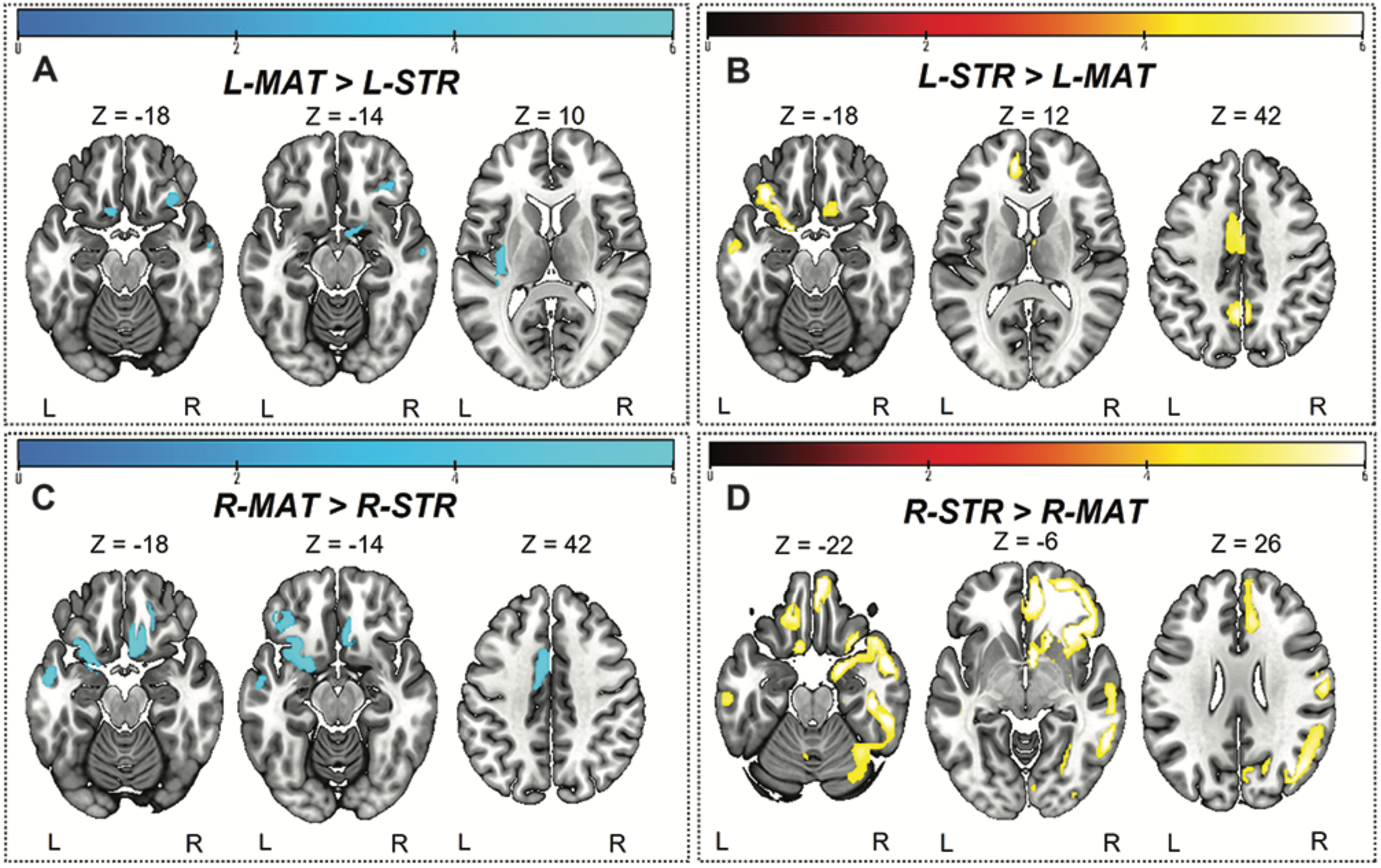
Significant differences between striosome- and matrix-seeded networks in the (a) left-matrix (L-MAT), (b) left-striosome (L-STR), (c) right-matrix (R-MAT), and (d) right-striosome (R-STR). Higher t-values indicate higher connectivity. Images are displayed in anatomical convention, where the left hemisphere is shown on the left side of the image.

**Figure 3:**
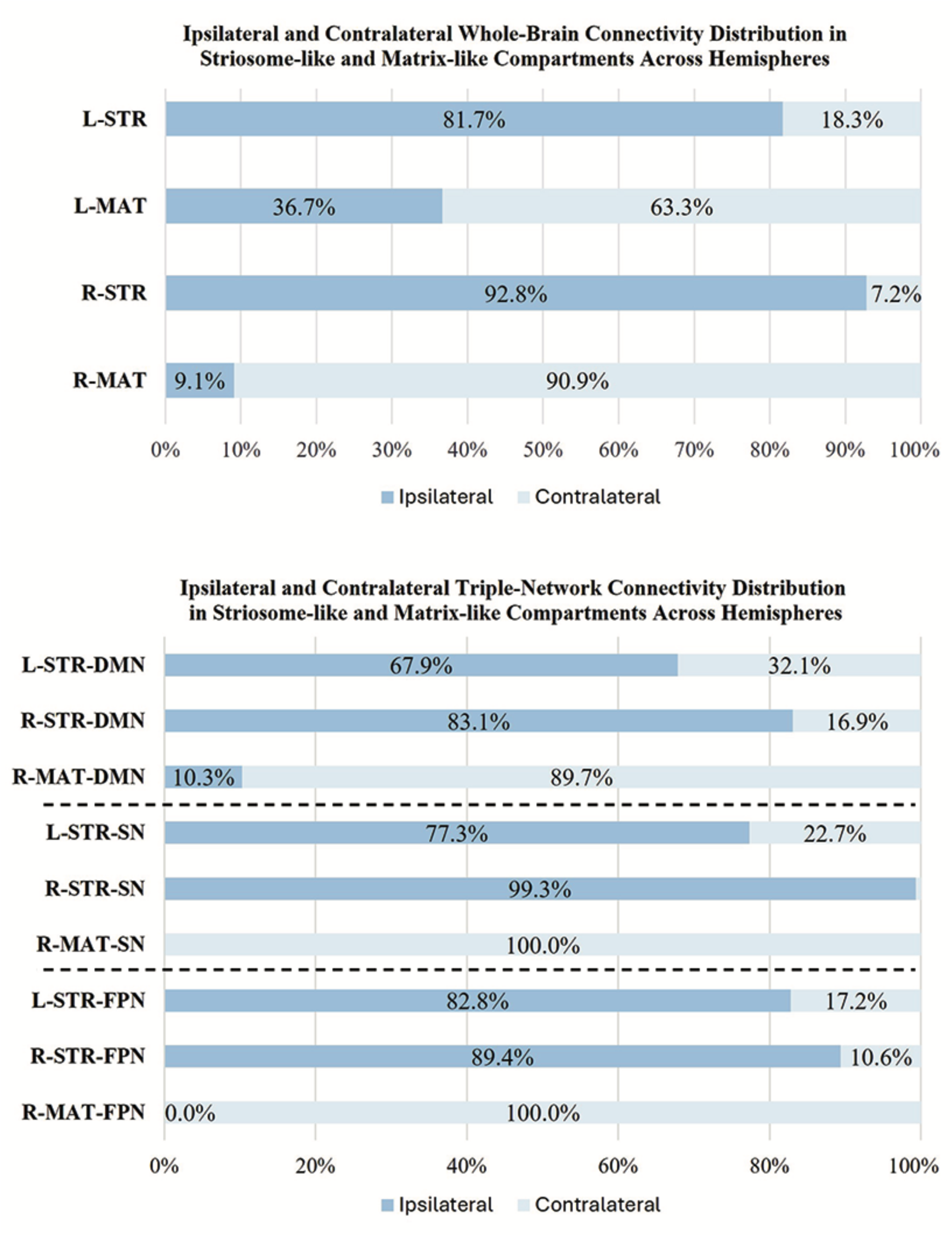
Comparison of ipsilateral and contralateral connectivity patterns in striosome and matrix compartments across hemispheres. The figure shows the distribution of whole-brain (top panel) and triple-network (bottom panel: DMN, default mode network; SN, salience network; FPN, fronto-parietal network) connectivity for ipsilateral and contralateral connections within striosome- and matrix-like compartments. Note that the L-MAT had no significant contrasts for DMN, SN, or FPN, so these comparisons are not shown in this figure.

The L-STR also demonstrated a high degree of ipsilateral connectivity, at 81.7%, though less biased than the R-STR. The L-MAT shows a more balanced distribution, with 63.3% of its significant voxels in the ipsilateral and 36.7% in the contralateral hemisphere, indicating moderate inter-hemispheric communication. Overall, the R-STR and R-MAT regions showed starkly contrasting hemispheric connectivity patterns, while the left hemisphere compartment-like seeds showed the same ipsilateral:contralateral pattern, but with smaller hemispheric bias.

In addition to compartment-specific hemispheric differences in rsFC, we identified rsFC differences in distinct regions in both hemispheres. L-MAT maps showed higher functional connectivity in brain regions involved in sensory-emotional integration and auditory processing, such as the left superior orbitofrontal cortex (sup-OFC), posterior insula (PIns), Heschl’s gyrus (HES), and the right inferior orbitofrontal cortex (Inf-OFC). In contrast, L-STR showed connectivity in relatively few right hemisphere regions, but these were associated with the DMN (precuneus (PCUN), sup-OFC, posterior cingulate cortex [PCC]). The L-STR also showed higher rsFC to several left hemisphere regions that contribute to cognitive-emotional integration, social cognition, and executive control (middle temporal gyrus (MTG), PCUN, middle and inferior OFC [mid-OFC,sup-OFC], anterior insula (AIns), and middle and anterior cingulate cortex [MCC, ACC]).

Compartment-specific differences between right hemisphere seeds were similar to those in the left hemisphere but were more robust. R-MAT maps showed higher functional connectivity in regions in the left hemisphere: temporal gyrus (TG), middle and superior temporal pole (MTP, STP), OFC, amygdala (AMYG), fusiform (FUS), AIns, postcentral gyrus, MCC, ACC, and PCUN. R-MAT also showed greater connectivity in regions in the right hemisphere: PIns and sup-OFC. In contrast, R-STR demonstrated substantially stronger connectivity, compared to R-MAT, in several right hemisphere regions, including the TG, MTP, and STP, inferior and middle frontal gyri (IFG, MFG) and occipital gyrus regions, including the middle and inferior occipital gyrus (MOG, IOG), cuneus (CUN), AIns, MCC, ACC, angular gyrus (ANG), gyrus rectus, hippocampus (HIPP), para-HIPP, AMYG, lingual (LING), FUS, and several cerebellar subregions (lobules VI, VII, VIII, IX; crus I, crus II). In the left hemisphere, R-STR connectivity was significantly elevated in the superior and middle frontal gyrus (SFG, MFG), ACC, postcentral gyrus, and the region encompassing both the gyrus rectus and the medial orbitofrontal cortex (mOFC), a key area involved in olfactory processing.

R-MAT and R-STR are connected to some of the same regions in opposite hemispheres, reflecting a lateralized, potentially opposing, pattern of functional connectivity. The left TG, MTP, STP, AMYG, FUS, and AIns had significant connectivity with R-MAT, while these same regions in the right hemisphere had significant connectivity with R-STR. This similarity in connectivity suggests that, despite their distinct compartmental roles, R-MAT and R-STR influence parallel networks on opposite sides of the brain, highlighting a unique cross-hemispheric mirroring of their connectivity profiles.

### 3.3 Influence of Precise Voxel Location on Striatal Compartmentalization

We have proposed that selecting striatal voxels based on biases in their structural connectivity can replicate the anatomic features of striosome and matrix that were demonstrated through histology. However, an alternate explanation for these biases in connectivity is that cortico-striatum projections are somatotopically organized, independent of compartment-level organization, and our striosome-like and matrix-like voxels are simply reflecting the connectivity biases of their local environment. We set out to determine if our precisely selected voxels, or the neighborhood in which they were embedded, was the primary driver of differences in rsFC. We jittered the location of our striosome-like and matrix-like seed voxels by ±0-3 voxels, at random and independently in each plane. The average root-mean-square distance shifted by individual voxels was modest: 2.9 voxels for randomized-striosome, 3.1 voxels for randomized-matrix. This measure incorporated the absolute value of shifted distance. The average distance shifted in each plane (which included both positive and negative shifts) was minimal: 0.37 voxels (range: 0.07-1.2 voxels). As planned, shifting voxels at random led to small but appreciable changes in location for individual voxels, but no meaningful change in the averaged location of whole striatal masks; on average, jittered masks occupied the same “neighborhood” as our precise striosome-like and matrix-like masks.

These location-shifted seed masks had no differences in rsFC; comparing functional correlations between striosome-shifted and matrix-shifted seeds, there were no significant voxels. This is consistent with our prior findings in structural connectivity in a separate diffusion MRI dataset (Funk, Hassan et al. 2023): small shifts in voxel location are sufficient to erase the compartment-specific biases in connectivity. These results suggest that the rsFC biases we observed are highly dependent on the precise localization of voxels within the striatum, rather than the broader neighborhood in which they reside.

### 3.4 Default Mode network (DMN)

We observed substantial, widespread compartment-specific differences in rsFC with the DMN (Figure 4). For L-STR seeds, most significant clusters (67.9%) were within the left hemisphere. Similarly, for R-STR seeds, 83.1% of significant clusters were within the right hemisphere. In contrast, the R-MAT seeds had 89.7% of their significant clusters in the contralateral hemisphere. There were no significant differences for connectivity between DMN and L-MAT. These DMN-specific interactions follow the same fundamental pattern demonstrated for whole-brain networks: striosome-like rsFC is predominantly ipsilateral, while matrix-like rsFC is less robust and predominantly contralateral.

**Figure 4:**
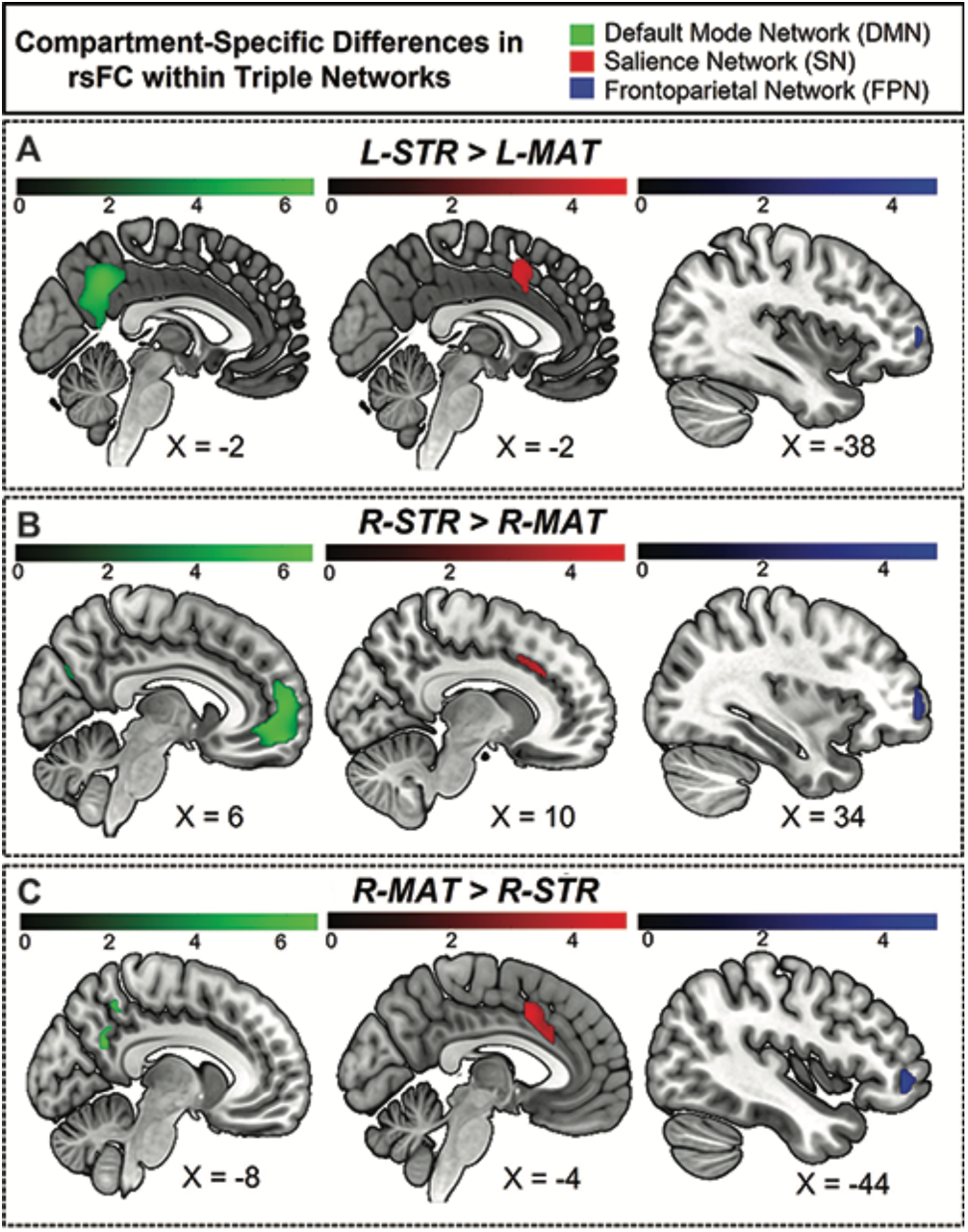
Hemisphere-specific distribution of striosome- and matrix-like compartment rsFC in the triple network. Panel A illustrates the regions where the L-STR exhibits stronger functional connectivity than the L-MAT compartment. Panel B illustrates the regions where the R-STR has stronger connectivity than the R-MAT. Panel C illustrates the regions where the R-MAT has stronger connectivity than the R-STR. Note that the L-MAT > L-STR contrast yielded no significant differences, and therefore is not pictured here. Green indicates regions within the DMN, red indicates regions in the SN, and blue indicates regions in the FPN. The color bars correspond to t-values, with higher values reflecting stronger connectivity.

The volume of significant clusters underscores the dominance of striosome-like seeds in influencing the DMN. Significant correlations between L-STR and DMN had a volume of 9,656 mm^3^, while the L-MAT to DMN correlations had a volume of zero. Correlations between R-STR and DMN were substantially larger (14,464 mm^3^) than R-MAT (1,008 mm^3^). Comparing the total number of significant voxels (both hemispheres) between the compartments, striosome-like seeds were 24-fold more likely to correlate with the DMN than matrix-like seeds.

The L-STR dominated connectivity in the precuneus of both hemispheres; L-MAT had no significant correlations. R-STR connectivity was evident in left prefrontal and right middle temporal regions, while R-MAT connectivity was observed in only a small area of the left medial precuneus (MNI coordinates: [-9.7, −53.3, 31.6]). Significant differences for L-STR-seeded maps were also noted in the ACC, CUN, PCC, and mPFC in the left hemisphere. R-STR-seeded connectivity was significantly higher in the mPFC, MTG, angular gyrus (ANG), CUN, PCUN, and ACC.

Regarding the overlap between hemispheric connectivity, the Dice Similarity Coefficient (DSC) between rsFC seeded by L-STR and R-STR ([L-STR > L-MAT] ∪ [R-STR > R-MAT]), was zero. Overlap between L-STR and R-MAT was minimal (DSC = 0.13). Intriguingly, the sole overlapping region between L-STR and R-MAT was the left precuneus, suggesting that this area may be a unique convergence point that links the otherwise distinct connectivity pattern of striosome-like and matrix-like seeds with the DMN.

### 3.5 Salience network (SN)

The L-STR and R-STR seeds demonstrated predominantly ipsilateral rsFC with SN nodes. Specifically, the L-STR seed exhibited 77.3% connectivity within the left hemisphere, while the R- STR seed showed 99.3% connectivity within the right hemisphere. Regions with significant L-STR– SN connectivity included the left MCC and ACC. The L-MAT showed no significant differences. R- STR–SN connectivity was significant in the right AIns, ACC, and SMG. R-MAT–SN connectivity was significant in the left ACC and AIns, continuing the pattern of contralateral connectivity for matrix. Indeed, 100% of the R-MAT–SN significant clusters were in the left hemisphere.

The volume of significant differences suggests that striosome-like seeds were dominant in influencing SN activity. The volume of significant correlations between L-STR and SN was 1,936 mm^3^. R-STR–SN correlation volume was 4,480 mm^3^, substantially larger than in the left hemisphere. In contrast, the volume of significant R-MAT–SN correlations was 2,888 mm^3^. Comparing the total number of significant voxels, striosome-like seeds were 2.2-fold more likely to influence the SN than matrix-like seeds.

Regarding the overlap in rsFC within the SN, the DSC between left and right striosome-like seeds ([L-STR > L-MAT] ∪ [R-STR > R-MAT]) was zero. However, L-STR and R-MAT showed substantial overlap (DSC = 0.51). The overlap between L-STR and R-MAT was restricted to the MCC, suggesting that this area may be a unique convergence point that links the distinct connectivity patterns of striosome-like and matrix-like seeds with the SN.

### 3.6 Frontoparietal network (FPN)

The ipsilateral dominance for striosome-like seeds and contralateral dominance for matrix-like seeds was also evident in the FPN. For both L-STR and R-STR, connectivity with the FPN was largely ipsilateral (82.8% and 89.4% of significant connectivity, respectively). FPN regions with significant connectivity to striosome-like seeds (both left and right) included the MFG and IFG. In contrast, no significant differences were found for L-MAT. R-MAT–FPN connectivity was entirely contralateral (left MFG and IFG).

The volume of significant differences indicates that striosome-like seeds had a stronger influence on FPN activity than matrix-like seeds. The volume of significant correlations between L-STR and the FPN was 512 mm³, whereas no significant correlations were observed in L-MAT–FPN connectivity. In contrast, the R-STR–FPN correlations had a volume of 2272 mm³, while the R-MAT–FPN correlations had a volume of 792 mm³. In combined hemispheres, striosome-like seeds were 3.5-fold more likely to influence the FPN than matrix-like seeds. The DSC for FPN regions was lower than for the DMN or SN: zero for L-STR︶R-STR, and minimal for L-STR︶R-MAT (DSC = 0.16). This overlap was restricted to the left IFG. While this overlap suggests that the left IFG may link connectivity between striosome-like and matrix-like networks, the substantially smaller overlap makes this association less clear for FPN than for regions in the DMN and SN.

### 3.7 N-1 Analysis: Validating Findings in the Anterior Insula

Among our ten compartment-favoring bait regions, the anterior insula was unique in also being an ROI for subsequent rsFC assessments. To ensure that our previously-noted rsFC findings in the anterior insula were accurate (not distorted by its use as a bait region), we carried out a second striatal parcellation that left out the anterior insula: an N-1 parcellation. We then used these N-1 striosome-like and matrix-like seeds as the seeds for rsFC with the insula and compared these network maps to our original connectivity maps.

Our N-1 analysis was not meaningfully different from our original parcellation (Figure 5). The differences in cluster volume (N-1 vs. original) were minimal, ranging from a 0.26% decrease to a 0.43% increase. The overlap of N-1 and original clusters was high. The DSC (N-1 vs. original) for L- MAT had moderate overlap of 0.61, while the R-MAT exhibited a higher DSC (0.85). Overlap within networks seeded by striosome-like voxels was high, with L-STR DSC of 0.75, and R-STR DSC of 0.89. Since the volume and location of compartment-specific significant clusters changed very little when parcellated without the anterior insula, we conclude that these compartment-specific patterns in rsFC were not distorted by the use of the anterior insula as a bait region for striatal parcellation.

**Figure 5:**
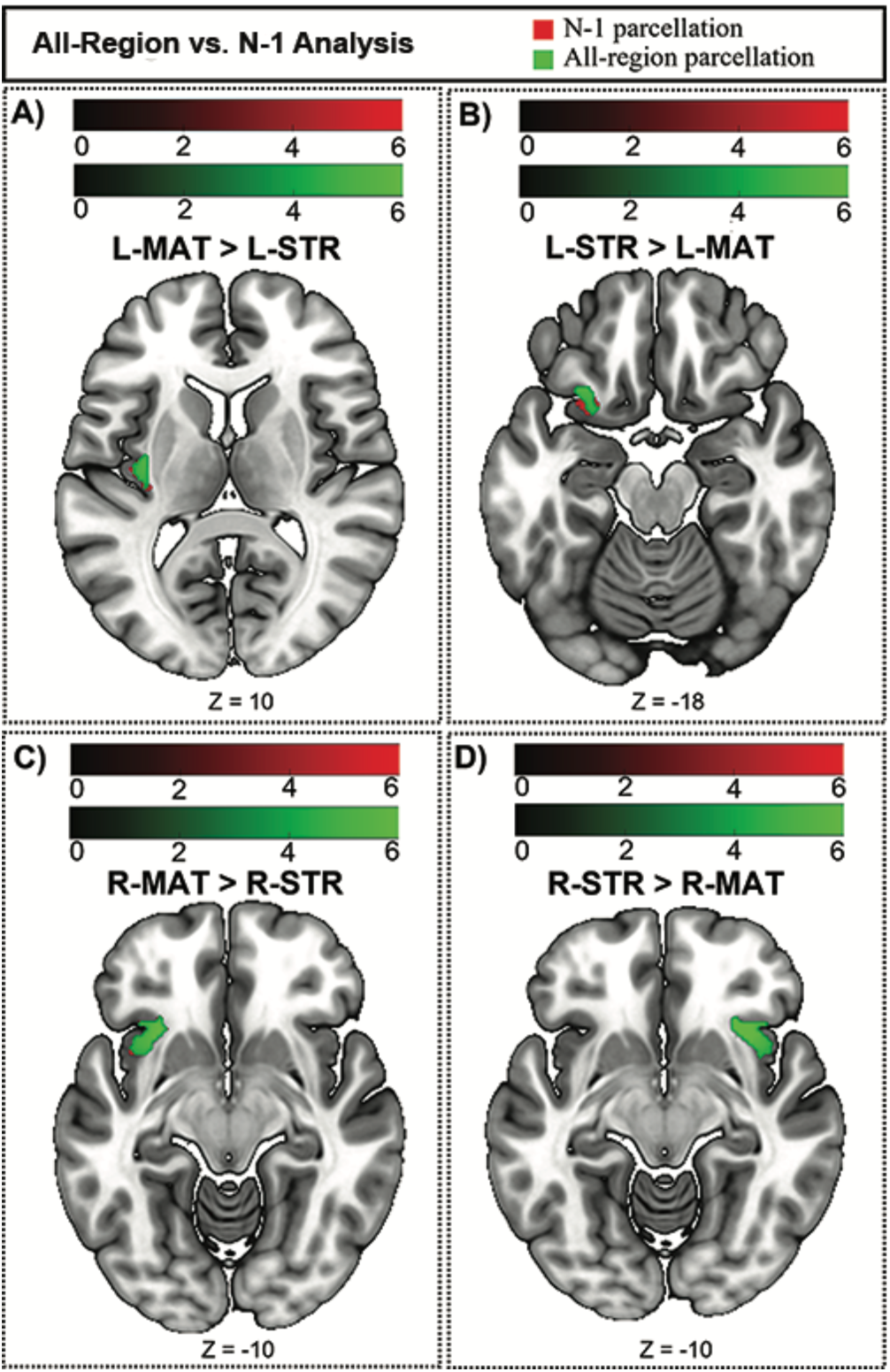
Compartment-specific bias in functional connectivity could be influenced by the use of the anterior insula as a bait region and subsequently as a site for whole-brain and salience network connectivity. Green voxels (original, all-region parcellation) are displayed in front of red voxels (N-1 parcellation). Connectivity is changed very little by leaving out the anterior insula. The panel (A) shows the regions where the L-MAT exhibits stronger functional connectivity than the L-STR compartment. The panel (B) illustrates the regions where the L-STR has stronger connectivity than the L-MAT. The panel (C) shows the regions where the R-MAT has stronger connectivity than the R-STR. The panel (D) shows the regions where the R-STR has stronger connectivity than the R-MAT. The color bars correspond to t-values, with higher values reflecting stronger connectivity. Images are displayed in anatomical convention, where the left hemisphere is shown on the left side of the image.

## 4 Discussion

For decades, the embryologic, pharmacologic, hodologic, and spatial segregation of striosome and matrix have suggested that the compartments’ functions are also distinct (Graybiel and Ragsdale Jr 1978; Brimblecombe and Cragg 2017; Prager, Dorman et al. 2020; Reiner, Chehimi et al. 2024). Intriguing behavioral assessments in non-human primates (Canales and Graybiel 2000; Karunakaran, Amemori et al. 2021) and rodents (Smith and Graybiel 2014; Friedman, Homma et al. 2015; Friedman, Homma et al. 2017) are consistent with this hypothesis: striosome-specific activation is essential for mood- and stress-influenced learning and decision making. Reward appears to be a striosome-specific function as well, since electrical self-stimulation is reinforcing when electrodes are placed in striosome, but not in matrix (White and Hiroi 1998). These striosome-specific functions are a small fraction of the tasks in which the striatum is involved, and do not attempt to map the implications of compartment-selective injury in human neuropsychiatric diseases (Crittenden and Graybiel 2011). In this study, we began to explore the differences in rsFC between voxels with striosome-like and matrix-like patterns of structural connectivity, and mapped rsFC with both whole-brain and function-specific (triple) networks. Though we have previously demonstrated that striosome-like and matrix-like voxels are embedded in distinct cortico-striato-thalamo-cortical <u>structural</u> networks (Funk, Hassan et al. 2023; Funk, Hassan and Waugh 2024), to the best of our knowledge this is the first demonstration that the compartments occupy distinct <u>functional</u> networks. Our findings suggest that compartment-specific functions in humans may be mediated by the influence of striosome or matrix acting within segregated functional networks. Moreover, these compartment-network relationships suggest that disease-specific symptoms may arise from selective decoupling of striosome or matrix from their segregated functional networks. While these insights into the rsFC of the striatal compartments are promising, our study has several limitations that must be considered.

First, the current method depends on the identification of striosome-like and matrix-like voxels using differential structural connectivity (probabilistic tractography). The accuracy of our rsFC findings is contingent on how well these compartment-like seed masks represent striosome and matrix at the tissue level. While we have previously demonstrated that striatal parcellation has a test-retest error rate of just 0.14% (Waugh, Hassan et al. 2022), and tractography-based parcellation replicates the patterns of compartment-biased structural connectivity demonstrated through injected tract tracers in animals for cortical and subcortical regions (Waugh, Hassan et al. 2022; Funk, Hassan et al. 2023; Funk, Hassan and Waugh 2024), our inferrential method is not the equivalent of histologic identification of striosome and matrix. The resolution of diffusion MRI is another important limitation of this method. The 1.25 mm isotropic voxels used here match the maximum diameter of the human striosome (Graybiel and Ragsdale 1978; Holt, Graybiel and Saper 1997), but smaller and obliquely sampled striosome branches are certain to produce partial volume effects and loss of discrimination between the compartments. At this and larger resolutions, every striosome-like voxel will include some fraction of matrix tissue. Inaccurate parcellation due to limited resolution could lead to imprecise seed voxels and yield incorrect functional connectivity maps. Although the limitations of resolution must be considered, the present study has a more precise resolution than our previous characterizations of compartment-like connectivity – where we demonstrated, in each of five distinct MRI datasets, that striosome-like and matrix-like voxels follow the expected relative abundance, spatial distribution, and biased structural connectivity demonstrated in animal and human histology (Waugh, Hassan et al. 2022; Funk, Hassan et al. 2023; Funk, Hassan and Waugh 2024; Marecek, Krajca et al. 2024).

Second, this study has the potential limitation of relying on predefined brain networks rather than defining functional networks *de novo* directly from the current dataset. While predefined brain networks, such as those identified in standard atlases or by previous studies, offer a convenient reference framework, they may not capture the unique network configurations present in the current dataset. Such individualized networks may be especially important for studying the striatal compartments in populations with neuropsychiatric diseases or neurodevelopmental disorders.

We observed that striosome-like seeds had markedly larger clusters of significant rsFC with the whole-brain and within the triple-network, compared to matrix-like seeds. Notably, striosome-like rsFC dominated even though the two compartment-like masks had equal volume and their distribution within the striatum was largely adjacent to the opposing compartment. These findings were highly dependent on the precise locations of these compartment-like voxels, as shifting their location by a few millimeters was sufficient to eliminate all rsFC. This heightened connectivity may be attributable to the intrinsic properties of striosome MSNs, which have higher input resistance and greater depolarization than matrix MSNs (Miura, Saino-Saito et al. 2007), as well as lower thresholds for firing action potentials and higher firing frequencies (Crittenden, Lacey et al. 2017). These properties suggest that striosome MSNs could be more readily excitable than matrix MSNs, facilitating their stronger influence on functional brain networks. However, further research is needed to confirm these findings and to identify mechanisms underlying these differences.

Previous studies investigating the striatal compartments have largely focused on animal models to explore aspects such as spatial distribution (Kubota and Kawaguchi 1993), neurochemical composition (Holt, Graybiel and Saper 1997), structural connectivity (Ragsdale Jr and Graybiel 1991), cortical organization (Ragsdale Jr and Graybiel 1990), functional segregation (Rajakumar, Elisevich and Flumerfelt 1993), and differences in dopamine regulation (Perreault, Hasbi et al. 2011; Prager and Plotkin 2019; McGregor, McKinsey et al. 2019). While these studies provided essential insights into compartment-specific differences, understanding the specific roles of striosome and matrix in human brain function and their involvement in diseases requires investigations using human subjects. Human histological studies, both in non-diseased and diseased brains, have begun to yield insights into the unique roles the compartments may play in neuropsychiatric conditions (e.g., studies examining human brain tissue for striatal compartment pathology in disorders like Huntington disease or Parkinson disease; Vonsattel, Myers et al. 1985; Reiner, Albin et al. 1988; Stephen,Kathleen and Oleh 1988; Gibb and Lees 1991). However, studies of post-mortem tissue cannot investigate the dynamic functional properties of human brain networks.

The striatal compartments are functionally separable and play distinct roles in the brain’s processing of information and regulation of behaviors (Saka and Graybiel 2003). One hypothesis proposes a functional differentiation between striosome and matrix MSNs, with the former being more involved in motivational aspects of behavioral function and the latter being more involved in sensory-motor functioning (Grabiel 1990; Eblen and Graybiel 1995; Graybiel 1997). The striosome is involved in integrating limbic (emotional and motivational) information, modulating dopaminergic activity, and influencing reward-related behaviors. In contrast, the matrix primarily performs sensorimotor integration and motor control, receiving the majority of inputs from sensorimotor areas and regulating movement (Crittenden and Graybiel 2011; Brimblecombe and Cragg 2017; McGregor, McKinsey et al. 2019). These functional distinctions have largely been inferred from patterns of connectivity in animal studies. The findings of the current study provide evidence for widespread, compartment-specific differences in rsFC in healthy humans, affecting a range of functional networks involved in cognitive processing, decision-making, self-referential thinking, and coordinating responses to internal and external events – the DMN, SN, and FPN, respectively (Scolari, Seidl-Rathkopf and Kastner 2015; Wen, Mitchell and Duncan 2020; Schimmelpfennig, Topczewski et al. 2023).

Hemispheric lateralization in the human brain is evident in key cognitive and motor tasks (Hervé, Zago et al. 2013) and is reflected in the ventral striatum as well, where the left and right hemispheres have distinct functional connectivity patterns (Zhang, Hu et al. 2017). Zhang et al. found that connectivity with the left striatum was primarily to regions involved in self-control and internal processes, while the right striatum connected more with areas related to attention and external actions. In the present study, we found that the right striatum had larger compartment-specific rsFC biases than the left, for each of the networks we assessed. Likewise, we observed a robust distinction between the ipsilateral dominance of striosome-like connectivity and the contralateral dominance of matrix-like connectivity. This pattern was consistent across whole-brain and task-specific networks (DMN, SN, and FPN). These findings suggest a compartment-specific role in hemispheric processing, where striosomes may support the refinement of motor and cognitive processes on the same side of the body, possibly contributing to more localized, fine-tuned control (Fujiyama, Unzai and Karube 2019). In contrast, our results suggest that the matrix compartment facilitates inter-hemispheric communication, coordinating broader, bilateral movements and processes that require integration across hemispheres.

Our functional connectivity analyses revealed significant region-specific commonalities and differences, particularly in core triple-network nodes. Key regions such as the OFC, AIns, MCC, anterior ACC, and PCUN exhibited consistent connectivity differences between left and right, and between compartment-like networks (L-STR, L-MAT, R-STR, R-MAT), underscoring their critical roles in modulating cognitive control, emotional regulation, and self-referential processing within the triple-network (Botvinick, Cohen and Carter 2004; Rolls 2004; Cavanna and Trimble 2006; Buckner, Andrews-Hanna and Schacter 2008; Craig 2009; Shackman, Salomons et al. 2011; Uddin 2015).

These areas are crucial nodes in the interaction between the DMN, SN, and FPN, suggesting that striosome and matrix compartments may differentially influence the balance between these networks, depending on the hemisphere and compartment involved. For instance, L-STR showed extensive connectivity in the left hemisphere, particularly in regions like the MTG, PCUN, AIns, MCC, and ACC. These areas are heavily involved in integrating sensory input with higher-order cognitive processes, such as decision-making and emotional responses, which are key functions of the SN and DMN. In contrast, L-MAT was associated with more limited regions, such as the sup-OFC, PIns, and HES in the left hemisphere, and the Inf-OFC in the right hemisphere. L-STR was connected with a broad array of regions while L-MAT had more restricted connectivity. R-STR demonstrated broad connectivity in the right hemisphere, including regions like the temporal gyrus (TG), MTP, STP, FG, and occipital regions (middle and inferior occipital gyri), along with the amygdala and cerebellum (lobule: VI, VII, VIII, IX, crus I, crus II), while also including left hemisphere regions like the SFG and MFG. The involvement of regions such as the TG, MTP, STP, FG, and occipital cortices, regions associated with auditory processing, visual processing, memory, and higher-order cognitive functions (Cabeza and Nyberg 2000; Kanwisher, McDermott and Chun 2002; Grill-Spector and Malach 2004; Hickok and Poeppel 2007; Duncan 2010). This suggests that R-STR may have a role in multimodal and associative sensory processing, while primary sensory processing is restricted to matrix.

The inclusion of left hemisphere regions in R-STR’s connectivity map indicates that it may play a role in cross-hemispheric regulation. This contralateral connectivity could be significant for tasks that require coordination between the hemispheres. Functional connectivity with the cerebellum suggests that R-STR might also influence motor, cognitive, or emotional tasks that require modeling and revision based on sensory and social feedback (Schmahmann 2021; Zhang,Duan et al. 2023). On the other hand, R-MAT is notable for its extensive connectivity in the left hemisphere, encompassing regions such as TG, MTP, STP, IFG, AMYG, and FUS, with significant right hemisphere connectivity observed in the PIns and sup-OFC, supporting the lateralization of matrix-like networks, where R- MAT preferentially engaged in interhemispheric rsFC and had distinct patterns of rsFC in each hemisphere. This lateralization may underpin specialized functions in sensory, emotional, and cognitive processing. These findings highlight the nuanced and compartment-specific functional connectivity with the striatum, reflecting both shared and unique contributions to brain network dynamics.

Striosome-like connectivity dominated the triple networks (DMN, SN, and FPN), particularly in the right hemisphere. This was particularly true in the DMN, which is primarily involved in self-referential thinking, episodic memory, and cognitive-sensory integration (Dixon, Moodie et al. 2022). The lateralization of DMN connectivity may suggest that in the right hemisphere, the striosome plays a critical role in integrating spatial and sensory information, potentially enhancing visuospatial processing and social cognition. In the SN, which filters and prioritizes salient stimuli (Uddin 2015), the more extensive right hemisphere striosome connectivity with the AIns and ACC underscores its potential contribution to emotional regulation (Coen, Yágüez et al. 2009; Stevens, Hurley and Taber 2011) and cognitive control (Sterzer and Kleinschmidt 2010). This hemispheric bias could imply a specialization of the striosome for processing emotional and sensory information within the SN. Finally, in the FPN, which governs executive functions and decision-making, the ipsilateral dominance of striosome connectivity, particularly in frontal cortices, suggests that it may play a key role in lateralized cognitive processes such as sequential reasoning and motor planning in the left hemisphere (Corballis 2014), while the right hemisphere supports visuospatial tasks (Hartikainen 2021). These findings point to a lateralized, compartment-specific modulation of higher-order cognitive functions across the triple networks, with the right striosome exerting a broader influence on network dynamics than matrix.

Interestingly, though striosome-like and matrix-like rsFC was markedly divergent by hemisphere, our most- and least-biased networks (L-STR and R-MAT) partially overlapped (DSC = 0.23). This overlap suggests that despite the distinct functional connectivity patterns typically observed between striosome-like and matrix-like networks, there are limited areas where left and right hemisphere networks may impinge upon the other compartment. The regions of overlap (all left hemisphere) include the anterior insula, anterior and middle cingulate cortices, precuneus, and orbitofrontal cortex. These regions are key nodes in the salience and default mode networks that are critical for emotional processing, cognitive control, and self-referential thinking (Dixon, Moodie et al. 2022; Schimmelpfennig, Topczewski et al. 2023). This convergence may reflect a level of integration between the striosome and matrix compartments, where they jointly contribute to key cognitive and emotional processes. The shared connectivity in these areas could be essential for coordinating complex behaviors that require the integration of emotional, cognitive, and self-referential information, underscoring the central role of compartment-specific functional networks.

In conclusion, this study provides strong evidence that in humans, the striosome and matrix compartments operate within segregated functional networks. Each compartment’s functional networks are organized differently (ipsilateral vs. contralateral; global vs. local). Striosome-like voxels primarily show ipsilateral connectivity and appear to be involved in hemisphere-specific processing. In contrast, matrix-like voxels exhibit stronger contralateral connectivity, indicating a more global role that integrates information across hemispheres. These patterns of connectivity suggest that striosome may be more specialized for local processing within the same hemisphere, whereas matrix compartments may contribute to broader, cross-hemispheric network dynamics. This distinction between ‘global’ and ‘local’ network organization helps to further elucidate the differential roles the striatal compartments play in brain function. Striosome-like seeds exhibited widespread functional connectivity with key nodes of the default mode, salience, and frontoparietal networks, suggesting involvement of the striosome in higher-order cognitive functions, emotional regulation, and integrative processing. In contrast, matrix-like seeds demonstrated more limited connectivity within these networks, potentially reflecting its specialized role in sensorimotor integration and motor control (Deng, Lanciego et al. 2015; Graybiel and Matsushima 2023). Task-based fMRI could be utilized to assess the activation of compartment-like seeds during sensory, motor, limbic, and executive tasks. While the temporal pattern of activation with tasks is an essential part of understanding the function of striosome and matrix, such investigations are beyond the scope of the current study.

Each striatal compartment is embedded in segregated intrinsic functional networks, providing an anatomic substrate for striosome and matrix to have distinct functional roles. The functional architecture of the striatum is intricate and highly localized – slight shifts in the location of striatal seeds eliminated all rsFC – suggesting that particular functions may also localize to limited zones within striosome or matrix. Future compartment-specific interventions may also require this level of anatomic precision. Linking specific symptoms with compartment-specific alterations in functional connectivity may suggest novel neuropathological mechanisms underlying neuropsychiatric disorders associated with striatal dysfunction, such as Parkinson disease (Albin, Young and Penney 1989), Huntington disease (Rosenblatt and Leroi 2000), or obsessive-compulsive disorder (Burguiere, Monteiro et al. 2015). Mapping the contributions of each striatal compartment to specific functional networks is essential for identifying the functions of striosome and matrix.

## Data availability statement

Publicly available datasets were analyzed in this study. This data can be found here: https://www.humanconnectome.org/study/hcp-young-adult/document/1200-subjects-data-release The code, bait, seed, and exclusion masks necessary to complete striatal parcellation can be accessed here: github.com/jeff-waugh/Striatal-Connectivity-based-Parcellation.

## Author Contributions

AS: data analysis, initial manuscript drafting, and critical revision of the manuscript. AF: acquisition of data and critical manuscript revision. JW: data acquisition, analysis, and interpretation, initial manuscript drafting, and critical manuscript revision. All authors contributed to the article and approved the final version for submission.

## References

Albin, R. L., A. B. Young and J. B. Penney (1989). “The functional anatomy of basal ganglia disorders.” Trends in neurosciences 12(10): 366–375.

Barto, A. G. (1995). “Adaptive critics and the basal ganglia.”

Binnewijzend, M. A., M. M. Schoonheim, E. Sanz-Arigita, A. M. Wink, W. M. van der Flier, N. Tolboom, S. M. Adriaanse, J. S. Damoiseaux, P. Scheltens and B. N. van Berckel (2012). “Resting-state fMRI changes in Alzheimer’s disease and mild cognitive impairment.” Neurobiology of aging 33(9): 2018–2028.

Botvinick, M. M., J. D. Cohen and C. S. Carter (2004). “Conflict monitoring and anterior cingulate cortex: an update.” Trends in cognitive sciences 8(12): 539–546.

Bressler, S. L. and V. Menon (2010). “Large-scale brain networks in cognition: emerging methods and principles.” Trends in cognitive sciences 14(6): 277–290.

Brimblecombe, K. R. and S. J. Cragg (2017). “The striosome and matrix compartments of the striatum: a path through the labyrinth from neurochemistry toward function.” ACS chemical neuroscience 8(2): 235–242.

Buckner, R. L., J. R. Andrews-Hanna and D. L. Schacter (2008). “The brain’s default network: anatomy, function, and relevance to disease.” Annals of the new York Academy of Sciences 1124(1): 1–38.

Burguiere, E., P. Monteiro, L. Mallet, G. Feng and A. M. Graybiel (2015). “Striatal circuits, habits, and implications for obsessive–compulsive disorder.” Current opinion in neurobiology 30: 59–65.

Burke, R. and K. Baimbridge (1993). “Relative loss of the striatal striosome compartment, defined by calbindin-D28k immunostaining, following developmental hypoxic-ischemic injury.” Neuroscience 56(2): 305–315.

Cabeza, R. and L. Nyberg (2000). “Imaging cognition II: An empirical review of 275 PET and fMRI studies.” Journal of cognitive neuroscience 12(1): 1–47.

Canales, J. J. and A. M. Graybiel (2000). “A measure of striatal function predicts motor stereotypy.” Nature neuroscience 3(4): 377–383.

Cavanna, A. E. and M. R. Trimble (2006). “The precuneus: a review of its functional anatomy and behavioural correlates.” Brain 129(3): 564–583.

Chakravarty, M. M., J. L. Rapoport, J. N. Giedd, A. Raznahan, P. Shaw, D. L. Collins, J. P. Lerch and N. Gogtay (2015). “Striatal shape abnormalities as novel neurodevelopmental endophenotypes in schizophrenia: a longitudinal study.” Human brain mapping 36(4): 1458–1469.

Coen, S. J., L. Yágüez, Q. Aziz, M. T. Mitterschiffthaler, M. Brammer, S. C. Williams and L. J. Gregory (2009). “Negative mood affects brain processing of visceral sensation.” Gastroenterology 137(1): 253–261. e252.

Corballis, M. C. (2014). “Left brain, right brain: facts and fantasies.” PLoS biology 12(1): e1001767.

Craig, A. D. (2009). “How do you feel—now? The anterior insula and human awareness.” Nature reviews neuroscience 10(1): 59–70.

Crittenden, J. R. and A. M. Graybiel (2011). “Basal Ganglia disorders associated with imbalances in the striatal striosome and matrix compartments.” Frontiers in neuroanatomy 5: 59.

Crittenden, J. R., C. J. Lacey, F.-J. Weng, C. E. Garrison, D. J. Gibson, Y. Lin and A. M. Graybiel (2017). “Striatal cholinergic interneurons modulate spike-timing in striosomes and matrix by an amphetamine-sensitive mechanism.” Frontiers in Neuroanatomy 11: 20.

Crittenden, J. R., P. W. Tillberg, M. H. Riad, Y. Shima, C. R. Gerfen, J. Curry, D. E. Housman, S. B. Nelson, E. S. Boyden and A. M. Graybiel (2016). “Striosome–dendron bouquets highlight a unique striatonigral circuit targeting dopamine-containing neurons.” Proceedings of the National Academy of Sciences 113(40): 11318–11323.

Deng, Y., J. Lanciego, L. K.-L. Goff, P. Coulon, P. Salin, P. Kachidian, W. Lei, N. Del Mar and A. Reiner (2015). “Differential organization of cortical inputs to striatal projection neurons of the matrix compartment in rats.” Frontiers in systems neuroscience 9: 51.

Desban, M., C. Gauchy, M. Kemel, M. Besson and J. Glowinski (1989). “Three-dimensional organization of the striosomal compartment and patchy distribution of striatonigral projections in the matrix of the cat caudate nucleus.” Neuroscience 29(3): 551–566.

Desban, M., M. Kemel, J. Glowinski and C. Gauchy (1993). “Spatial organization of patch and matrix compartments in the rat striatum.” Neuroscience 57(3): 661–671.

Dixon, M. L., C. A. Moodie, P. R. Goldin, N. Farb, R. G. Heimberg, J. Zhang and J. J. Gross (2022). “Frontoparietal and default mode network contributions to self-referential processing in social anxiety disorder.” Cognitive, Affective, & Behavioral Neuroscience: 1–12.

Donoghue, J. P. and M. Herkenham (1986). “Neostriatal projections from individual cortical fields conform to histochemically distinct striatal compartments in the rat.” Brain research 365(2): 397–403.

Duncan, J. (2010). “The multiple-demand (MD) system of the primate brain: mental programs for intelligent behaviour.” Trends in cognitive sciences 14(4): 172–179.

Eblen, F. and A. M. Graybiel (1995). “Highly restricted origin of prefrontal cortical inputs to striosomes in the macaque monkey.” Journal of Neuroscience 15(9): 5999–6013.

Flaherty, A. W. and A. M. Graybiel (1993). “Two input systems for body representations in the primate striatal matrix: experimental evidence in the squirrel monkey.” Journal of Neuroscience 13(3): 1120–1137.

Friedman, A., D. Homma, B. Bloem, L. G. Gibb, K.-i. Amemori, D. Hu, S. Delcasso, T. F. Truong, J. Yang and A. S. Hood (2017). “Chronic stress alters striosome-circuit dynamics, leading to aberrant decision-making.” Cell 171(5): 1191–1205. e1128.

Friedman, A., D. Homma, L. G. Gibb, K.-i. Amemori, S. J. Rubin, A. S. Hood, M. H. Riad and A. M. Graybiel (2015). “A corticostriatal path targeting striosomes controls decision-making under conflict.” Cell 161(6): 1320–1333.

Friedman, A., E. Hueske, S. M. Drammis, S. E. T. Arana, E. D. Nelson, C. W. Carter, S. Delcasso, R. X. Rodriguez, H. Lutwak and K. S. DiMarco (2020). “Striosomes mediate value-based learning vulnerable in age and a Huntington’s disease model.” Cell 183(4): 918–934. e949.

Fujiyama, F., T. Unzai and F. Karube (2019). “Thalamostriatal projections and striosome-matrix compartments.” Neurochemistry international 125: 67–73.

Funk, A. T., A. A. Hassan, N. Brüggemann, N. Sharma, H. C. Breiter, A. J. Blood and J. L. Waugh (2023). “In humans, striato-pallido-thalamic projections are largely segregated by their origin in either the striosome-like or matrix-like compartments.” Frontiers in neuroscience 17: 1178473.

Funk, A. T., A. A. Hassan and J. L. Waugh (2024). “In humans, insulo-striate structural connectivity is largely biased toward either striosome-like or matrix-like striatal compartments.” bioRxiv.

Gerfen, C. R. (1984). “The neostriatal mosaic: compartmentalization of corticostriatal input and striatonigral output systems.” Nature 311(5985): 461–464.

Gibb, W. and A. Lees (1991). “Anatomy, pigmentation, ventral and dorsal subpopulations of the substantia nigra, and differential cell death in Parkinson’s disease.” Journal of Neurology, Neurosurgery & Psychiatry 54(5): 388–396.

Gimenez-Amaya, J. and A. Graybiel (1990). “Compartmental origins of the striatopallidal projection in the primate.” Neuroscience 34(1): 111–126.

Goldman-Rakic, P. S. (1982). “Cytoarchitectonic heterogeneity of the primate neostriatum: subdivision into island and matrix cellular compartments.” Journal of Comparative Neurology 205(4): 398–413.

Grabiel, A. (1990). “Neurotransmitter and neuromodulator in the basal ganglia.”

Graybiel, A. M. (1997). “The basal ganglia and cognitive pattern generators.” Schizophrenia bulletin 23(3): 459–469.

Graybiel, A. M. and S. T. Grafton (2015). “The striatum: where skills and habits meet.” Cold Spring Harbor perspectives in biology 7(8): a021691.

Graybiel, A. M. and A. Matsushima (2023). “Striosomes and matrisomes: scaffolds for dynamic coupling of volition and action.” Annual review of neuroscience 46(1): 359–380.

Graybiel, A. M. and C. W. Ragsdale, Jr. (1978). “Histochemically distinct compartments in the striatum of human, monkeys, and cat demonstrated by acetylthiocholinesterase staining.” Proc Natl Acad Sci U S A 75(11): 5723–5726.

Graybiel, A. M. and C. W. Ragsdale Jr (1978). “Histochemically distinct compartments in the striatum of human, monkeys, and cat demonstrated by acetylthiocholinesterase staining.” Proceedings of the National Academy of Sciences 75(11): 5723–5726.

Grill-Spector, K. and R. Malach (2004). “The human visual cortex.” Annu. Rev. Neurosci. 27(1): 649–677.

Haber, S. N., K.-S. Kim, P. Mailly and R. Calzavara (2006). “Reward-related cortical inputs define a large striatal region in primates that interface with associative cortical connections, providing a substrate for incentive-based learning.” Journal of Neuroscience 26(32): 8368–8376.

Hartikainen, K. M. (2021). “Emotion-attention interaction in the right hemisphere.” Brain sciences 11(8): 1006.

Hedreen, J. C., S. Berretta and C. L. White III (2024). “Postmortem neuropathology in early Huntington disease.” Journal of Neuropathology & Experimental Neurology 83(5): 294–306.

Hervé, P.-Y., L. Zago, L. Petit, B. Mazoyer and N. Tzourio-Mazoyer (2013). “Revisiting human hemispheric specialization with neuroimaging.” Trends in cognitive sciences 17(2): 69–80.

Hickok, G. and D. Poeppel (2007). “The cortical organization of speech processing.” Nature reviews neuroscience 8(5): 393–402.

Holt, D. J., A. M. Graybiel and C. B. Saper (1997). “Neurochemical architecture of the human striatum.” Journal of Comparative Neurology 384(1): 1–25.

Holt, D. J., A. M. Graybiel and C. B. Saper (1997). “Neurochemical architecture of the human striatum.” J Comp Neurol 384(1): 1–25.

Houk, J., J. Adams and A. Barto (1995). “A Model of How the Basal Ganglia Generate and Use Neural Signals that Predict Reinforcement, Models of Information Processing in the Basal Ganglia (eds. JC Houk, JL Davis and DG Beiser), 249/270.” Preprint at.

Johnston, J. G., C. R. Gerfen, S. N. Haber and D. van der Kooy (1990). “Mechanisms of striatal pattern formation: conservation of mammalian compartmentalization.” Developmental Brain Research 57(1): 93–102.

Jung, W. H., J. H. Jang, J. W. Park, E. Kim, E.-H. Goo, O.-S. Im and J. S. Kwon (2014). “Unravelling the intrinsic functional organization of the human striatum: a parcellation and connectivity study based on resting-state FMRI.” PloS one 9(9): e106768.

Kanwisher, N., J. McDermott and M. M. Chun (2002). “The fusiform face area: a module in human extrastriate cortex specialized for face perception.”

Karunakaran, K. B., S. Amemori, N. Balakrishnan, M. K. Ganapathiraju and K.-i. Amemori (2021). “Generalized and social anxiety disorder interactomes show distinctive overlaps with striosome and matrix interactomes.” Scientific Reports 11(1): 18392.

Kubota, Y. and Y. Kawaguchi (1993). “Spatial distributions of chemically identified intrinsic neurons in relation to patch and matrix compartments of rat neostriatum.” Journal of Comparative Neurology 332(4): 499–513.

Kuo, H.-Y. and F.-C. Liu (2020). “Pathological alterations in striatal compartments in the human brain of autism spectrum disorder.” Molecular brain 13: 1–4.

Lewis, M. and S.-J. Kim (2009). “The pathophysiology of restricted repetitive behavior.” Journal of neurodevelopmental disorders 1: 114–132.

Li, Y., H. Yao, P. Lin, L. Zheng, C. Li, B. Zhou, P. Wang, Z. Zhang, L. Wang and N. An (2017). “Frequency-dependent altered functional connections of default mode network in Alzheimer’s disease.” Frontiers in aging neuroscience 9: 259.

Marecek, S., T. Krajca, R. Krupicka, P. Sojka, J. Nepozitek, Z. Varga, C. Mala, J. Keller, J. Waugh and D. Zogala (2024). “Analysis of striatal connectivity corresponding to striosomes and matrix in de novo Parkinson’s disease and isolated REM behavior disorder.” npj Parkinson’s Disease 10(1): 124.

McGregor, M. M., G. L. McKinsey, A. E. Girasole, C. J. Bair-Marshall, J. L. Rubenstein and A. B. Nelson (2019). “Functionally distinct connectivity of developmentally targeted striosome neurons.” Cell reports 29(6): 1419–1428. e1415.

Menon, V. (2011). “Large-scale brain networks and psychopathology: a unifying triple network model.” Trends in cognitive sciences 15(10): 483–506.

Menon, V. and L. Q. Uddin (2010). “Saliency, switching, attention and control: a network model of insula function.” Brain structure and function 214: 655–667.

Mikula, S., S. K. Parrish, J. S. Trimmer and E. G. Jones (2009). “Complete 3D visualization of primate striosomes by KChIP1 immunostaining.” Journal of Comparative Neurology 514(5): 507–517.

Miura, M., S. Saino-Saito, M. Masuda, K. Kobayashi and T. Aosaki (2007). “Compartment-specific modulation of GABAergic synaptic transmission by μ-opioid receptor in the mouse striatum with green fluorescent protein-expressing dopamine islands.” Journal of Neuroscience 27(36): 9721–9728.

Nadel, J., S. Pawelko, J. Scott, R. McLaughlin, M. Fox, M. Ghanem, R. van der Merwe, N. Hollon, E. Ramsson and C. Howard (2021). “Optogenetic stimulation of striatal patches modifies habit formation and inhibits dopamine release.” Scientific Reports 11(1): 19847.

Perreault, M. L., A. Hasbi, B. F. O’Dowd and S. R. George (2011). “The dopamine D1–D2 receptor heteromer in striatal medium spiny neurons: evidence for a third distinct neuronal pathway in basal ganglia.” Frontiers in neuroanatomy 5: 31.

Prager, E. M., D. B. Dorman, Z. B. Hobel, J. M. Malgady, K. T. Blackwell and J. L. Plotkin (2020). “Dopamine oppositely modulates state transitions in striosome and matrix direct pathway striatal spiny neurons.” Neuron 108(6): 1091–1102. e1095.

Prager, E. M. and J. L. Plotkin (2019). “Compartmental function and modulation of the striatum.” Journal of neuroscience research 97(12): 1503–1514.

Ragsdale Jr, C. W. and A. M. Graybiel (1988). “Fibers from the basolateral nucleus of the amygdala selectively innervate striosomes in the caudate nucleus of the cat.” Journal of Comparative Neurology 269(4): 506–522.

Ragsdale Jr, C. W. and A. M. Graybiel (1990). “A simple ordering of neocortical areas established by the compartmental organization of their striatal projections.” Proceedings of the National Academy of Sciences 87(16): 6196–6199.

Ragsdale Jr, C. W. and A. M. Graybiel (1991). “Compartmental organization of the thalamostriatal connection in the cat.” Journal of Comparative Neurology 311(1): 134–167.

Rajakumar, N., K. Elisevich and B. Flumerfelt (1993). “Compartmental origin of the striato-entopeduncular projection in the rat.” Journal of Comparative Neurology 331(2): 286–296.

Reiner, A., R. L. Albin, K. D. Anderson, C. J. D’Amato, J. B. Penney and A. B. Young (1988). “Differential loss of striatal projection neurons in Huntington disease.” Proceedings of the National Academy of Sciences 85(15): 5733–5737.

Reiner, B. C., S. N. Chehimi, R. Merkel, S. Toikumo, W. H. Berrettini, H. R. Kranzler, S. Sanchez-Roige, R. L. Kember, H. D. Schmidt and R. C. Crist (2024). “A single-nucleus transcriptomic atlas of medium spiny neurons in the rat nucleus accumbens.” Scientific Reports 14(1): 18258.

Rolls, E. T. (2004). “The functions of the orbitofrontal cortex.” Brain and cognition 55(1): 11–29.

Rosenblatt, A. and I. Leroi (2000). “Neuropsychiatry of Huntington’s disease and other basal ganglia disorders.” Psychosomatics 41(1): 24–30.

Saka, E. and A. M. Graybiel (2003). “Pathophysiology of Tourette’s syndrome: striatal pathways revisited.” Brain and Development 25: S15–S19.

Salinas, A. G., M. I. Davis, D. M. Lovinger and Y. Mateo (2016). “Dopamine dynamics and cocaine sensitivity differ between striosome and matrix compartments of the striatum.” Neuropharmacology 108: 275–283.

Schimmelpfennig, J., J. Topczewski, W. Zajkowski and K. Jankowiak-Siuda (2023). “The role of the salience network in cognitive and affective deficits.” Frontiers in human neuroscience 17: 1133367.

Schmahmann, J. D. (2021). “Emotional disorders and the cerebellum: neurobiological substrates, neuropsychiatry, and therapeutic implications.” Handbook of clinical neurology 183: 109–154.

Schuetze, M., M. T. M. Park, I. Y. Cho, F. P. MacMaster, M. M. Chakravarty and S. L. Bray (2016). “Morphological alterations in the thalamus, striatum, and pallidum in autism spectrum disorder.” Neuropsychopharmacology 41(11): 2627–2637.

Scolari, M., K. N. Seidl-Rathkopf and S. Kastner (2015). “Functions of the human frontoparietal attention network: Evidence from neuroimaging.” Current opinion in behavioral sciences 1: 32–39.

Shackman, A. J., T. V. Salomons, H. A. Slagter, A. S. Fox, J. J. Winter and R. J. Davidson (2011). “The integration of negative affect, pain and cognitive control in the cingulate cortex.” Nature Reviews Neuroscience 12(3): 154–167.

Smith, K. S. and A. M. Graybiel (2014). “Investigating habits: strategies, technologies and models.” Frontiers in behavioral neuroscience 8: 39.

Stephen, J., S. Kathleen and H. Oleh (1988). “Uneven pattern of dopamine loss in the striatum of patients with idiopathic Parkinson’s disease.” New England J. Med 318: 876–880.

Sterzer, P. and A. Kleinschmidt (2010). “Anterior insula activations in perceptual paradigms: often observed but barely understood.” Brain Structure and Function 214(5): 611–622.

Stevens, F. L., R. A. Hurley and K. H. Taber (2011). “Anterior cingulate cortex: unique role in cognition and emotion.” The Journal of neuropsychiatry and clinical neurosciences 23(2): 121–125.

Uddin, L. Q. (2015). “Salience processing and insular cortical function and dysfunction.” Nature reviews neuroscience 16(1): 55–61.

Van Essen, D. C., S. M. Smith, D. M. Barch, T. E. Behrens, E. Yacoub, K. Ugurbil and W.-M. H. Consortium (2013). “The WU-Minn human connectome project: an overview.” Neuroimage 80: 62–79.

Van Essen, D. C., K. Ugurbil, E. Auerbach, D. Barch, T. E. Behrens, R. Bucholz, A. Chang, L. Chen, M. Corbetta and S. W. Curtiss (2012). “The Human Connectome Project: a data acquisition perspective.” Neuroimage 62(4): 2222–2231.

Vonsattel, J.-P., R. H. Myers, T. J. Stevens, R. J. Ferrante, E. D. Bird and E. P. Richardson Jr (1985). “Neuropathological classification of Huntington’s disease.” Journal of Neuropathology & Experimental Neurology 44(6): 559–577.

Waldvogel, H. J., D. Thu, V. Hogg, L. Tippett and R. L. Faull (2012). “Selective neurodegeneration, neuropathology and symptom profiles in Huntington’s disease.” Tandem Repeat Polymorphisms: Genetic Plasticity, Neural Diversity and Disease: 141–152.

Wang, K., K. Li and X. Niu (2021). “Altered functional connectivity in a triple-network model in autism with co-occurring attention deficit hyperactivity disorder.” Frontiers in psychiatry 12: 736755.

Waugh, J. L., A. A. Hassan, J. K. Kuster, J. M. Levenstein, S. K. Warfield, N. Makris, N. Brüggemann, N. Sharma, H. C. Breiter and A. J. Blood (2022). “An MRI method for parcellating the human striatum into matrix and striosome compartments in vivo.” Neuroimage 246: 118714.

Wei, M., J. Qin, R. Yan, K. Bi, C. Liu, Z. Yao and Q. Lu (2015). “Association of resting-state network dysfunction with their dynamics of inter-network interactions in depression.” Journal of affective disorders 174: 527–534.

Wen, T., D. J. Mitchell and J. Duncan (2020). “The functional convergence and heterogeneity of social, episodic, and self-referential thought in the default mode network.” Cerebral cortex 30(11): 5915–5929.

White, N. M. and N. Hiroi (1998). “Preferential localization of self-stimulation sites in striosomes/patches in the rat striatum.” Proceedings of the National Academy of Sciences 95(11): 6486–6491.

Whitfield-Gabrieli, S. and A. Nieto-Castanon (2012). “Conn: a functional connectivity toolbox for correlated and anticorrelated brain networks.” Brain connectivity 2(3): 125–141.

Xiao, X., H. Deng, A. Furlan, T. Yang, X. Zhang, G.-R. Hwang, J. Tucciarone, P. Wu, M. He and R. Palaniswamy (2020). “A genetically defined compartmentalized striatal direct pathway for negative reinforcement.” Cell 183(1): 211–227. e220.

Zhang, P., L. Duan, Y. Ou, Q. Ling, L. Cao, H. Qian, J. Zhang, J. Wang and X. Yuan (2023). “The cerebellum and cognitive neural networks.” Frontiers in human neuroscience 17: 1197459.

Zhang, S., S. Hu, H. H. Chao and C.-s. R. Li (2017). “Hemispheric lateralization of resting-state functional connectivity of the ventral striatum: an exploratory study.” Brain Structure and Function 222: 2573–2583.

Zhong, Y., L. Huang, S. Cai, Y. Zhang, K. M. von Deneen, A. Ren, J. Ren and A. s. D. N. Initiative (2014). “Altered effective connectivity patterns of the default mode network in Alzheimer’s disease: an fMRI study.” Neuroscience letters 578: 171–175.

